# Foraging strategies under extreme events: Contrasting adaptations by benthic macrofauna to drastic biogeochemical disturbance

**DOI:** 10.1101/2020.09.09.288985

**Authors:** Yiming V. Wang, Mario Lebrato, Li-Chun Tseng, Thomas Larsen, Nicolás Smith-Sánchez, Pei-Wen Lee, Juan-Carlos Molinero, Jiang-Shiou Hwang, Tin-Yam Chan, Dieter Garbe-Schönberg

## Abstract

Extreme events caused by global change are increasingly affecting the ocean’s biogeochemical cycling and ecosystem functioning, but it is challenging to observe how food webs respond to rapid habitat disturbances. Benthic communities are particularly vulnerable because their habitats are easily affected by extreme events. Here, we examined how benthic macrofauna responded to a “near shutdown” of shallow marine hydrothermal vents, triggered by M5.8 earthquake and C5 typhoon events. Despite reduced vent fluxes, we shows that the endemic vent crab *Xenograpsus testudinatus* continued to rely on chemosynthetic sulfur bacteria rather than photosynthetic sources. We posit this obligate nutritional dependence caused a population decline of vent crabs. In contrast, the non-endemic mollusks exhibited much greater dietary plasticity with no detectable impact on the population. Our study based on naturally occurring extreme events exemplifies how specialist species in marine system are particularly vulnerable to the unprecedented evolutionary and environmental pressures exerted by human activities worldwide.

## Introduction

Marine habitats and biogeochemical cycles are increasingly affected by extreme natural hazards and human disturbances, leading to rapid loss of biodiversity and altered food web functioning (*1*). Of all the oceanic habitats, benthic food webs are among the most vulnerable to these types of disturbances because they are often spatially fragmented and contain macrofauna assemblages that are dominated by rare, endangered, or endemic species (*2, 3*). Unfortunately, it is challenging to observe the trophodynamic effects of these large-scale and rapid environmental phenomena on benthic food webs first-hand because changes can occur suddenly making “before and after” monitoring unattainable (*4, 5*). Furthermore, these large scale environmental phenomenon cannot be easily imitated in mesocosm experiments (*6, 7*). One of the solutions to this challenge is to conduct longitudinal studies at easily accessible locations that are frequently affected by earthquakes, typhoons, and other extreme events. Shallow marine hydrothermal vents, also known as shallow vents, are ideal locations for longitudinal studies because their food webs are supported by both chemosynthetic and photosynthetic sources that are sensitive to extreme events causing changes in fluid flow, elemental cycling, and seawater mixing (*8-10*) (Supplementary text I).

Although ecologists have monitored food webs around shallow vents under relatively stable environmental conditions, no studies have yet examined how food webs respond to extreme events using data collected before and after drastic biogeochemical disruptions (*11-14*). To improve environmental management and protection of hydrothermal ecosystems, it is important understanding how vent benthic fauna respond to a transition from active to inactive, and how they rely on resources from surrounding non-chemosynthetic ecosystems (*15*). Furthermore, hydrothermal vent animals have been studied as model species to understand the origin of life on Earth as well as animal adaptation and evolution (*16*) because of their ability to use the geochemical energy source and tolerate high toxicity at the seafloor (*17*). Given that hydrothermal ecosystems are increasingly threatened by human activities and targeted for exploitation (*15, 18*), we are losing biodiversity and the opportunity to continue studying conditions similar to those analogous to environments that fostered the origin of life on Earth.

In 2016, one of the most heavily studied shallow marine vent systems in the world, Kueishantao Island (KST, 24.843°N, 121.951°E, site information is detailed Supplementary text II) off the coast of Taiwan, was hit by a M5.8 earthquake and a subsequent C5 typhoon within a few weeks (12^th^ May and 2^nd^-10^th^ July, respectively). These extreme events triggered underwater landslides, burying active vents and benthic habitats, causing a near “shutdown” of vent activity for almost two years (Fig. 1). The combined effect from the earthquake and typhoon decreased the toxicity and intensity of the discharging shallow vent fluids (*9*), the two most important factors for structuring KST’s food webs (*19, 20*). Before the extreme events, the active Yellow Vents (YV) and the semi-inactive White Vents (WV) at the KST had distinct biogeochemical properties that supported different food webs (Fig. 2). The high temperature and toxicity YV discharged gasses and fluids rich in CO_2_, sulfide (H_2_S) and sulfur dioxide (SO_2_) (*21*). In contrast, WV has low degassing activity and no discharge of toxic fluids (*21*). After the disturbance, the rapid change in the physical environment and reduced vent discharges led to dramatic shifts in the seawater’s biogeochemistry in both YV and WV (*9*). While dissolved inorganic carbon (DIC) and elements recovered to pre-disturbance concentrations after two year, previously abundant native yellow sulfur accretions near the active YV disappeared (Fig. 1 and 2). The diminished sulfur accretions could have led to a decrease in chemosynthetic sulfur bacterial biomass, and shifted the benthic food base from chemosynthetic to photosynthetic resources.

**Fig. 1.**
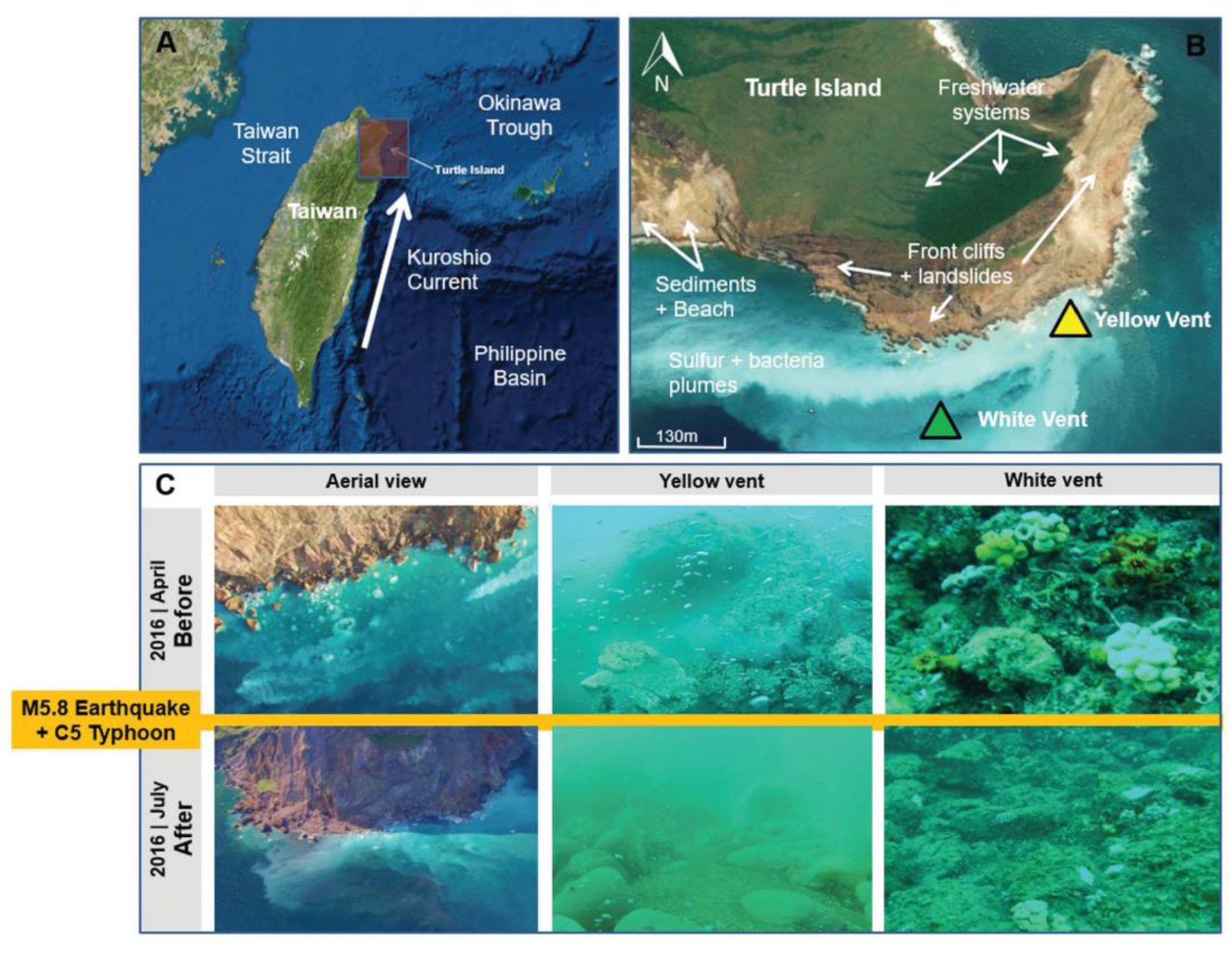
Kuishantao Island (Turtle Island, off Taiwan) geographical location and photographs showing the disturbance events that impacted shallow hydrothermal vent activity: (A) Shallow vents study area, located at a tectonic junction northeast of Taiwan and at the southern end of the Okinawa Trough. Its eastern side is directly hit by the Kuroshio Current. (B) Sampling locations at the Yellow Vent (YV), White Vent (WV), and old vent sites as well as Turtle Island east side details. (C) Aerial and submarine photographs showing the vent activity and seabed conditions at the sampling sites before and after the 5.8 Richter Magnitude earthquake on 12^th^ May 2016, and Category 5 typhoon Nepartak from 2^nd^ to 10^th^ of July 2016. Detailed time series aerial and submarine photos from 2011 to 2018 can be found in Lebrato et al. (2019). The databases are deposited at the NOAA National Center for Environmental Information (NCEI) under Accession Number 0175781 in https://data.nodc.noaa.gov/cgi-bin/iso?id=gov.noaa.nodc:0175781 with DOI:10.25921/6hy3-6d56.

**Figure 2.**
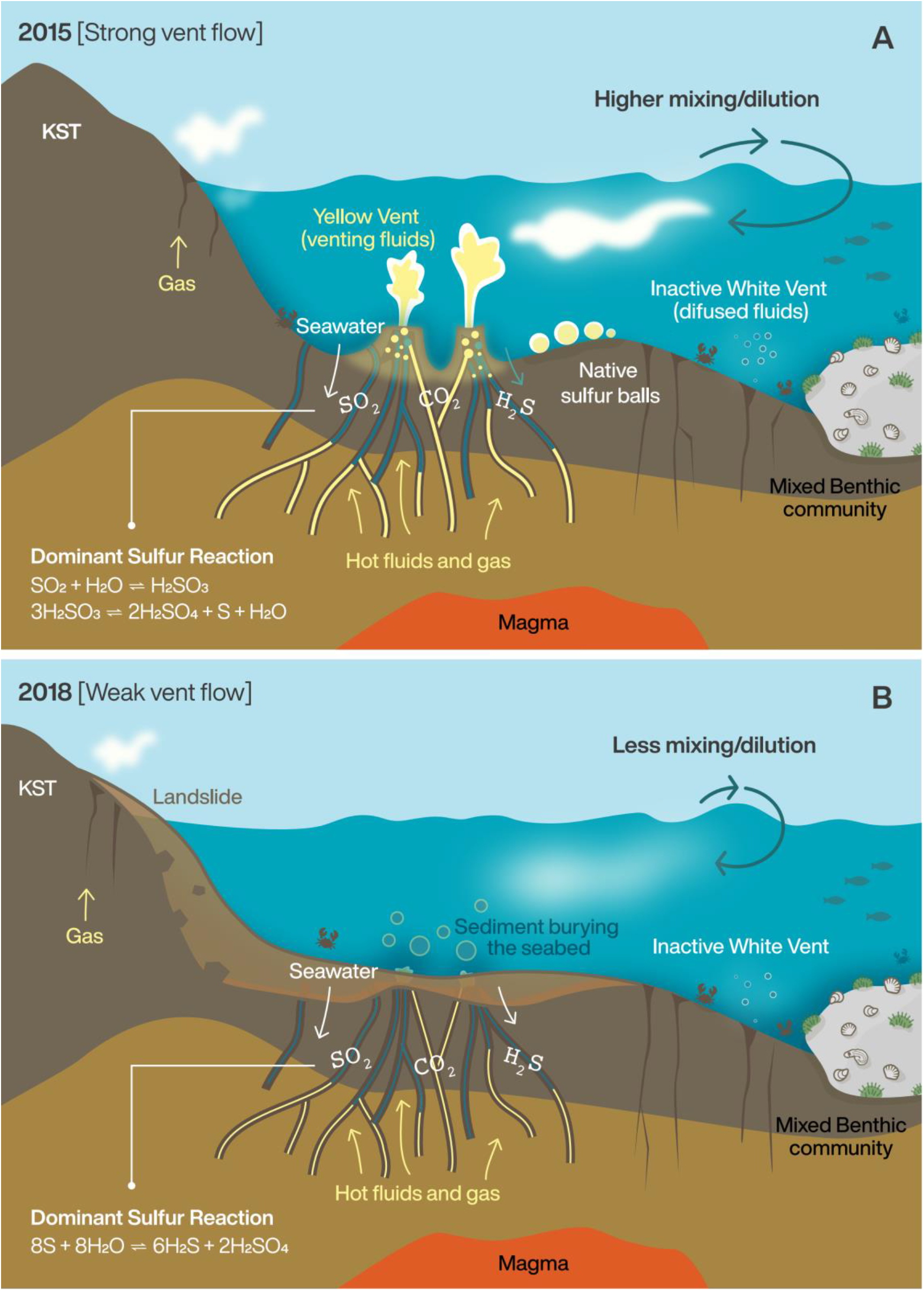
Diagrams showing dominant geochemical processes around Turtle Island shallow hydrothermal: A) 2015; B) 2017 and 2018. The geochemical changes owing to the Richter M5.8 earthquake and C5 Typhoon caused changes in sulfur element cycling. See text for detailed explanation. This figure is created by Michelle O’Reilly (Graphic Designer for the Max Planck Institute for the Science of Human History, Jena, Germany)

The only macrofaunal species (i.e. body length 0.5-50mm) to inhabit the hostile YV environment is the endemic vent crab *Xenograpsus testudinatus*. Its successful existence in the YV has been attributed to its physiological adaptation to highly acidic and toxic hydrothermal fluids (*19, 22, 23*). Although vent crabs are closely associated with toxic sulfur vent habitats, it is unclear whether they are obligate feeders of sulfur bacteria because vent crabs also rely on substantial contributions from photosynthetic derived sources by feeding on dead zooplankton, detritus and biofilms on vent rocks (*11-14*). Compared to YV, the less hostile WV has a much higher faunal diversity (Fig. 2). Besides the endemic vent crab, the WV hosts several generalist species such as mollusks capable of rapidly colonizing early succession habitats and using a variety of available resources (*19*). The coexistence of obligate and non-endemic vent species before and after the extreme events, makes KST an important site for understanding how individual members of benthic food webs are differentially affected by rapid and dramatic changes in biogeochemical and physical properties.

To understand how the shifts in biogeochemical cycles triggered by extreme events (2016) altered benthic food webs, we analyzed stable sulfur, carbon, and nitrogen isotope compositions (*δ*^34^S, *δ*^13^C, and *δ*^15^N) of the endemic vent crab (*X. testudinatus*), and three keystone mollusk species in the peripheral WV area (*19, 20*); the sessile gastropod *Thylacodes adamsii* (Mörch, 1859) and the two mobile gastropods, *Bostrycapulus gravispinosus* (Kuroda & Habe, 1950) and *Ergalatax contracta* (Reeve, 1846) in 2015 and 2018. These mollusks are considered keystone species in the vent area because they are pioneer species, abundant, and have a fairly high tolerance to the toxic conditions around these geothermal vents (*20*). The three mollusks represent different feeding strategies: *T. adamsii* and *B. gravispinosus* are filter feeders and *E. contracta* is a carnivore (*19, 20*). By combining ^34^S, a powerful tracer for tracking vent influence, with the classic dietary tracer ^13^C and trophic tracer ^15^N (*24-26*), we test the hypothesis that vent crabs have an obligate nutritional relationship with sulfur bacteria. In contrast, we posit that the dietary breadth of the generalist mollusk species will be much broader reflecting the dominant primary production resources at any given time.

## Results

### Changes in sulfur cycling at the KST before and after the disturbance

After these catastrophic events, the seabed landscape near the YV significantly changed; the new shoreline formed, the old vent mouths were buried, and the seabed was covered by debris and boulders from the landslide (Fig. 1 and 2). During the SCUBA diving expeditions of 2017 (July and August) and 2018 to the YV, we observed a dramatic decrease in venting intensities, the complete disappearance of sulfur ball deposits, and the collapse of the bright yellow sulfur chimneys. The sulfur deposits that were prevalent before the extreme events were either buried by landslide debris or dissolved by the changes in the vent’s fluid composition. By 2018, the sulfur chimney and deposit still did not recover. As illustrated in Figure 2A, when sulfur dioxide (SO_2_) predominated over hydrogen sulfide (H_2_S) in the vent fluids, the precipitation of native sulfur before the extreme events and the equilibrium between the different sulfur species can be explained as following equations (*27, 28*):

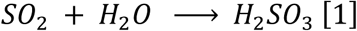

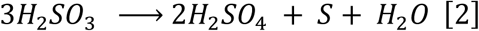

Under these oxidizing conditions, the production of sulfurous acid (H_2_SO_3_; reaction 1) resulted in elemental sulfur (reaction 2). After the disturbance (Fig. 2B), however, the conditions became more reductive, which lowered SO_2_ concentrations (*29*). Hence, the dominant equilibrium after extreme events can best be described as:

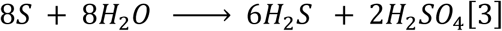

The absence of SO_2_ causes dissolution of native sulfur (*27*). Such a shift in sulfur species from a more oxidizing to a reductive state (decreasing ratio of SO_2_ to H_2_S) also resulted in a more pronounced ^34^S depletion of sulfides and the overall sulfur species in the vent fluids (*28, 30*).

### Interannual variability in stable isotope values of vent crabs between 2015 and 2018

We analyzed and compared the *δ*^13^C, *δ*^15^N, and *δ*^34^S of muscle tissues on vent crab (N=28) and found a large amount of variability and significant differences in isotope values for vent crabs (*X. testudinatus*) between 2015 and 2018 (Fig. 3, MANOVA Pillai’s Trace = 0.8, F_3,24_ = 34.9; *P* < 0.0001). However, the differences between the two years were mainly driven by changes in *δ*^34^S and *δ*^15^N (ANOVA, *P-adj* =0.003 and *P-adj* <0.0001 for *δ*^34^S *and δ*^15^N, respectively; Supplementary Table S2). Interestingly, the direction of change for *δ*^34^S and *δ*^15^N also differed, for example the average *δ*^34^S values decreased ∼2.4‰ whereas the *δ*^15^N values increased ∼2.1‰ from 2015 to 2018. In contrast, the *δ*^13^C values between these two years were not statistically different (ANOVA, *P-adj*=0.18, supplementary Table S2), and the mean *δ*^13^C values were 15.3 ±2.1‰ for 2015 and −16.2 ±1.1‰ for 2018.

**Figure 3.**
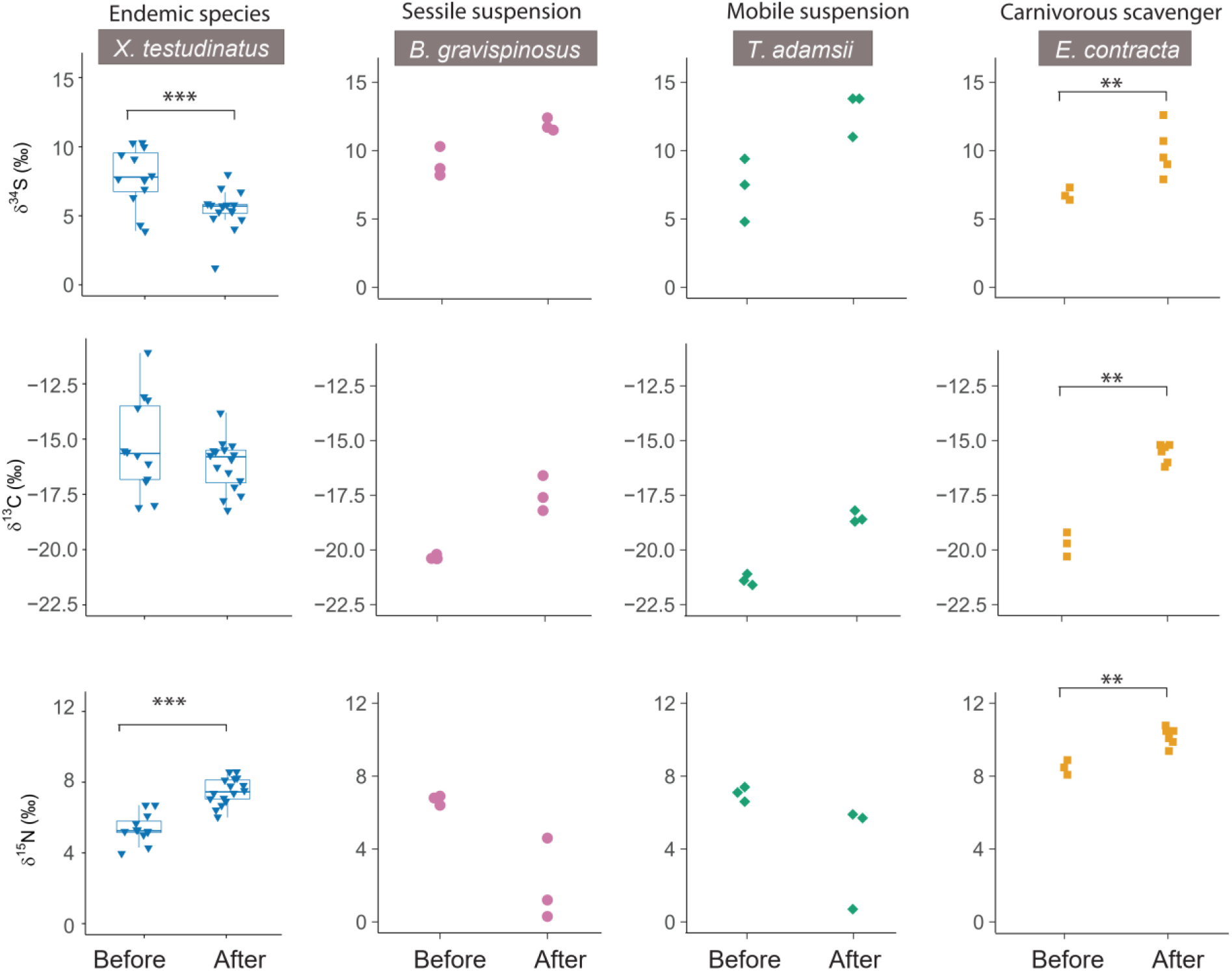
Inter-annual *δ*^34^S, *δ*^13^C and *δ*^15^N variability of benthic organism tissues in the Kuishantao Island in 2015 and 2018: vent crabs (*Xenograpsus testudinatus*); *Bostrycapulus gravispinosus*; *Thylacodes adamsii*; and *Ergalatax contracta* before and after the 2016 catastrophic events (Richter M5.8 earthquake and C5 Typhoon). *** denotes for *P*-adj <0.001 and ** denotes for *P*-adj <0.05.

### Interannual variability in δ^34^S, δ^13^C and δ^15^N of mollusks between 2015 and 2018

We observed apparent shifts in *δ*^34^S, *δ*^13^C, and *δ*^15^N for suspension feeding mollusks before and after extreme events although no statistical analyses were performed (due to small sample and effect size) (Fig. 3), with increases in *δ*^34^S (average of ∼3‰ and ∼5‰ for *B. gravispinosus* and *T. adamsii*, respectively), and *δ*^13^C values (average of ∼3‰ for both species). By contrast, *δ*^15^N values of these two suspension feeders decreased (∼5‰ for *B. gravispinosus* and ∼3‰ for *T. adamsii*) and their respective variability was much larger after these extreme events. For carnivorous mollusk *E. contracta*, there are significant increases in all the isotope values after the extreme events (two sample t-tests: t_4.8_=3.7, *P-adj*=0.02; t_7_=12.6 and *P-adj*<0.0001; t_6_=5.0 and *P-adj*=0.03 for *δ*^34^S, *δ*^13^C, and *δ*^15^N respectively). One striking difference between *E. contracta* and the two suspension feeders is the large variability between the *δ*^15^N values—their values greatly departed from each other by an average of 7‰ in 2018 compared to those in 2015 (Fig. 3A).

### Intra-annual isotope variability between vent crabs and mollusks in 2015 and 2018

We examined whether there was a significant difference in the intra-annual variability of isotope values (i.e. *δ*^13^C, *δ*^15^N, and *δ*^34^S) between vent crabs and the pooled mollusk species. We pooled all mollusks together because the vent crabs are considered the only endemic species in the KST hydrothermal vents. Our result show that the vent crabs and mollusks occupy different isotopic spaces but their difference are driven by a different isotope each year (Fig. 4). For example, the significant difference in 2015 (Pillai’s Trace = 0.80, F_3,17_ = 22.6; *P*-adj < 0.0001) was driven by differences in *δ*^13^C and *δ*^15^N values (ANOVA, *P-adj* <0.0001, supplementary Table S4), with *δ*^34^S not significantly differing between the two invertebrate groups (ANOVA, *P-adj* = 0.9). However, significant differences in 2018 (MANOVA Pillai’s Trace = 0.77, F_2,24_= 40.1; *P*-*adj* < 0.0001, Supplementary Table S5) were driven by changes in *δ*^34^S (ANOVA *P-adj* <0.0001, Table S5) instead of *δ*^13^C (*P*-*adj* =0.19). The average *δ*^13^C values were almost identical (−16.2‰ and −16.8‰ for vent crabs and mollusks, respectively) despite the large variations among macrofauna. We did not compared *δ*^15^N values in 2018 as the ranges of mollusks were wider than those of vent crabs.

**Figure 4.**
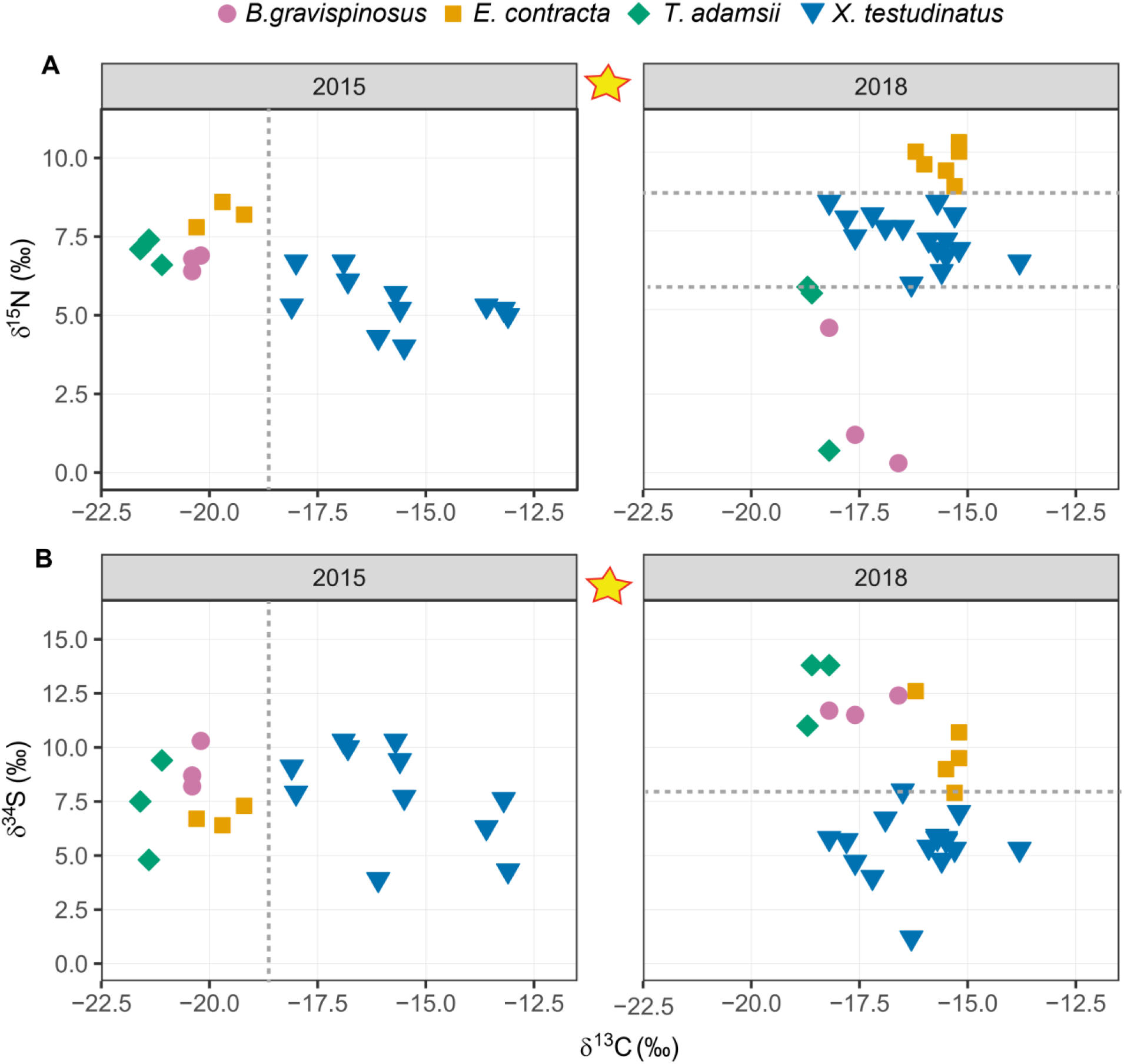
Paired stable isotope values of vent crabs (*Xenograpsus testudinatus*) and individual Molluscs (*Ergalatax contracta, Thylacodes adamsii, Bostrycapulus gravispinosus*) from the KST in years 2015, and 2018: A) *δ*^13^C and *δ*^15^N; B) *δ*^34^S and *δ*^13^C. The yellow stars denote Richter M5.8 earthquake and C5 Typhoon events in 2016.

### Stable isotope values of vent crabs by habitats and sex

No significant differences were detected between the vent crabs collected in the YV and WV habitats (MANOVA Pillai’s Trace = 0.49, F_3,12_= 3.80; *P*-*adj* = 0.08, Supplementary Table S6). The *δ*^34^S, *δ*^13^C and *δ*^15^N isotope signatures between the two habitats were identical (ANOVA *P*-*adj* =0.41, 0.03, and 0.16, respectively). Furthermore, no differences between sexes (female vs. male) in 2018 were found among vent crabs, despite some of the females carrying eggs at the time of sampling (Pillai’s Trace = 0.27, F_3,12_= 1.49; *P*-*adj* = 0.27, Supplementary Table S7). ANOVA analyses also show that *δ*^34^S, *δ*^13^C and *δ*^15^N between the two sexes are identical (*P*-*adj* =0.91, 0.26, and 0.53, respectively).

### Body sizes and population size changes before and after the extreme events

By comparing population data from 2004 and 2014 (*13, 19*) with our own underwater camera survey in 2018 (Supplementary Table S8), we found that the vent crab population at the YV decreased from average 23 specimens for every 25×25 cm^2^ in 2004 to average 11 and four specimens in 2014 and 2018, respectively. Despite lack of quantitative mollusk population data after the disturbances, we did not observe any apparent changes in mollusks’ population density after the earthquake.

To investigate whether the macrofanua species changed their body sizes in response to the biogeochemical changes, we also measured the sizes of vent crabs and mollusks except for the irregularly coiled *T. adamsii*. The size data are listed in the Supplementary Table S1. Two sample t-tests on body width on vent crabs and *E. contracta* before and after the extreme events showed that the body size did not change significantly after the disturbances (t_26_=-0.94, *P-adj* =0.35 for crabs; and t_7_=-0.141, *P-adj* =0.20 for *E. contracta*, respectively). The body size of *B. gravispinosus* is similar to previous observation (*20*) and also barely changed before and after the events (t_4_=-0.41, *P-adj* =0.05).

### Comparison of δ^34^S values between hydrothermal vent organisms and marine organisms in the open ocean

We compiled the *δ*^34^S data from this study with *δ*^34^S data and from the previously published literature to gain insight how *δ*^34^S values of the KST vent macrofauna compared to those deep-sea hydrothermal vent organisms as well as marine organisms in the open ocean (Fig. 5). Our data show endemic species vent crabs and non-endemic mollusks in the KST vents have similar *δ*^34^S ranges (average 9.2 and 7.7‰, respectively) before the extreme event. The *δ*^34^S values of both groups were between deep-sea vent fauna and pelagic fish or deep-sea benthic macrofauna (Fig. 5). The average *δ*^34^S values of both groups were lower than the middle point (10.5‰) of the two *δ*^34^S end members at the KST vent system: i) chemosynthetic food sources from KST native sulfur have a mean value of ∼1‰, and ii) photosynthetic sources of natural seawater sulfate has a mean value of ∼20‰ (*21*) (Fig. 5). After the extreme event, however, the *δ*^34^S values of vent crabs decreased towards values more typical of deep-sea hydrothermal benthic organisms (average of 5.1‰; Fig. 5), in spite of the diminished vent activities. In contrast, the *δ*^34^S values of mollusk species increased by an average of 7-8‰ and diverged from those of vent crabs by ∼12.3‰, thus resembling *δ* ^34^S values typical for pelagic fish and benthic organisms, ranging from 13‰ to 20‰ (Fig. 3B and Fig.5).

**Figure 5.**
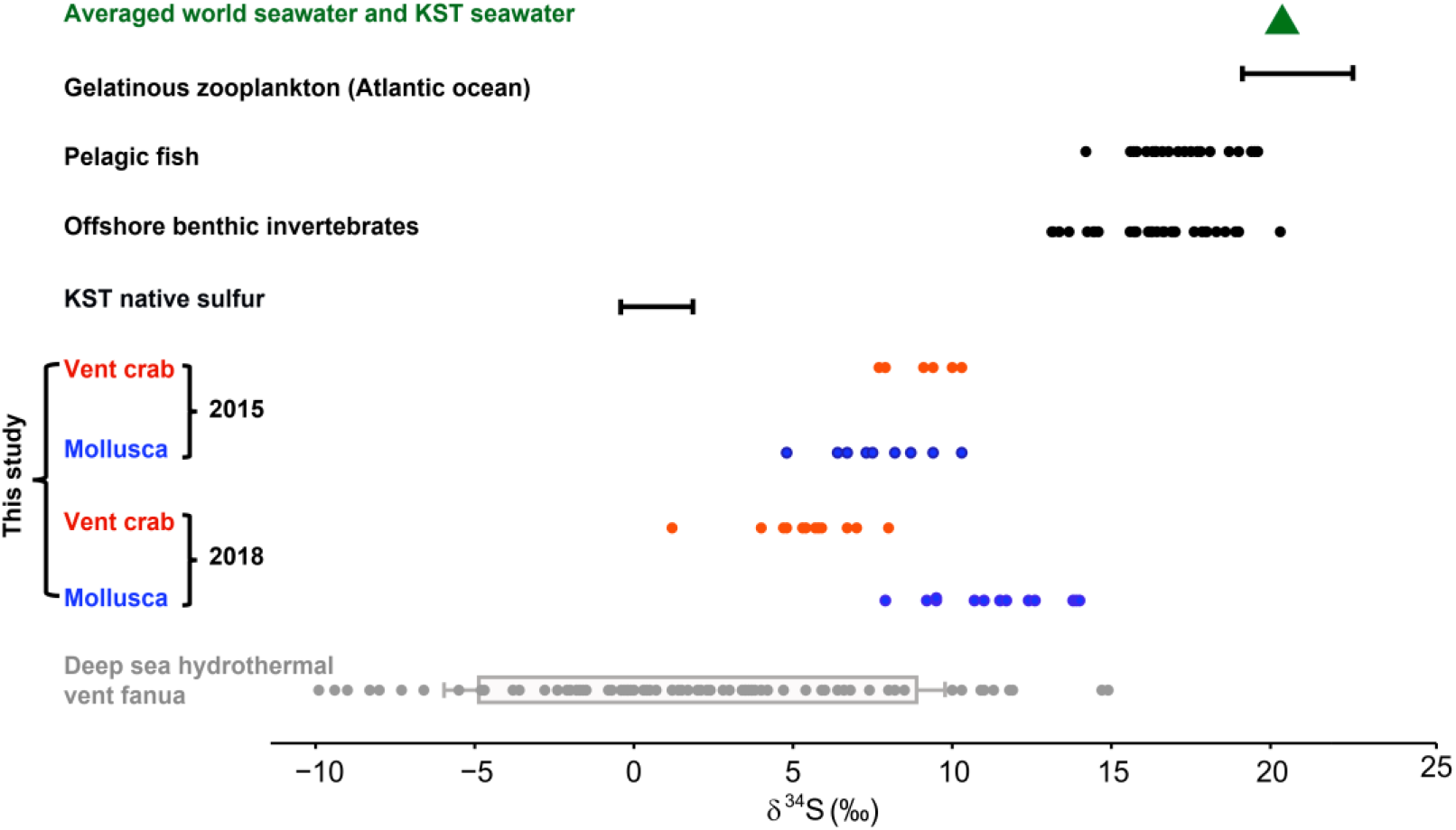
Compiled *δ*^34^S values of hydrothermal vent organisms and those of marine organisms in the open ocean: Filled circles with different colors indicates individual animal or tissues. The data based on previous published data are often mean values of several individual specimens of the same species. Filled triangle indicates the seawater sulfate from Gulf of Mexico and the KST. The *δ*^34^S value ranges are also shown for the native sulfur from the KST (*n*=5) and gelatinous zooplankton from the Atlantic Ocean (shelf seas) (*n*=675). Data reference: Jellyfish from the UK shelf (*42*); Pelagic fish (*43*); Offshore_benthic_invertebrates (*43-46*); KST native sulfur and seawater sulfate (*21*); Deep sea hydrothermal vent animals ((*45, 47, 48*)).

## Discussion

### Extreme events triggered drastic changes in sulfur cycling and isotope values

After the earthquake and typhoon in 2016, the major elements in the KST system experienced a period of rapid re-organization with a two-year near shutdown period and the disappearance of the previously abundant native yellow sulfur accretions on the seafloor near the active YV (Fig. 1 and 2). This suggests a major change in elemental sulfur cycling, namely a shift in sulfur speciation of KST fluids in the mixing zone between vent and sea waters where chemosynthetic bacteria uses H_2_S as an energy source (*31*). The shift in sulfur species from a more oxidizing to a reductive state (decreasing ratio of SO_2_ to H_2_S) also resulted in a more pronounced ^34^S depletion of sulfides and other sulfur species in the KST vent fluids. Indeed, studies from volcanic settings have shown that ^34^S of overall sulfur species become more depleted as a function of decreasing SO_2_/H_2_S (*27-30*). As a result, dietary sources associated with vents became more ^34^S depleted after these extreme events. Besides sulfur speciation, there was also a major reduction in sulfur fluid fluxes from the YV, leading to the increased influence of seawater (more ^34^S enriched) in the mixing zone. Taken together, these two sulfur cycling processes have a profound impact on the dietary sulfur availability of chemo- or photosynthetic sources for the vent benthic macrofauna (Fig. 5).

By using multiple isotopes, we were able to track how the KST food web responded to sudden biogeochemical changes. Due to the underlying biogeochemical complexity in the KST region, the source tracers ^34^S and ^13^C provided complementary dietary information. Under the strong venting activities prior to these extreme events, vent crabs and all mollusks from both the YV and WV showed consistent *δ*^34^S values but discordant *δ*^13^C values (Fig. 3). This result suggests that sulfur originating from the YV fluids are the largest sulfur source for both chemo- and photosynthetic production in the entire area, masking over the potential effects of the seawater’s sulfur source. Vent sulfur’s dominant influence in the vicinity of the KST area is not surprising because the S^2-^ concentration in the YV fluids was approximately five times higher than that of the WV fluids and 50 times of seawater in the Kuroshio Current (*21, 32*). Despite the much narrower range between chemo- and photosynthetic sources of ^13^C than ^34^S (*26, 33*), ^13^C is a more informative source tracer before the extreme events because of the small spatial difference in dissolved inorganic carbon (DIC) concentrations between the YV and the peripheral area (< 1.25) (*9*). After the extreme events disturbed the vent system, however, the reduced vent sulfur flux dampened the dominance of vent sulfur in the KST region, making ^34^S the most informative source tracer for chemo vs. photosynthetic origins (Fig. 5). For example, the divergence in *δ* ^34^S values between mollusks and vent crabs strongly suggest that the vent sulfur is no longer the dominant sulfur source for the entire KST area (Fig. 4). Our interpretation of the isotope data below is based on the underlying sulfur and carbon biogeochemistry in the KST region.

### Dietary response to the drastic biogeochemical changes and habitat disruption

The marked changes in sulfur cycling influenced the diet of hydrothermal macrofauna except for the endemic vent crab *X. testudinatus*. Before the disturbance, the more positive *δ*^13^C values in vent crabs compared to other vent invertebrates indicate that the crabs relied more on food derived from chemosynthetic sources than photosynthetic sources (Fig. 4). Our result corroborate previous trophic studies in the KST (*11, 14*). The consistently high *δ*^13^C values in vent crabs are attributed to the carbon intake from chemoautotrophic microbes that use high *δ*^13^C of DIC (average −18‰), and vent POM (∼17.4‰) (*11, 14*). Although DIC concentration peaked immediately after these extreme events, DIC values recovered to the same level as in May 2018 (*9*). The fact that *δ*^13^C values for vent crabs had similar ranges before and after the extreme events suggests that vent crabs continued to rely more on chemo-rather than photosynthetic dietary sources, despite the decreasing supply of vent derived dietary sources. Furthermore, the *δ*^34^S values in vent crab have become more negative after the disturbance affected the vents, indicating that vent crabs continue to obtain nutrients from the vent associated sources. However, the decrease in *δ*^34^S values was unlikely due to a proportional increase in the consumption of chemosynthetic sources (i.e. vent POM including sulfur based microbial biomass) as the vent activities decreased. Instead, the more negative *δ*^34^S values in vent crabs were likely driven by the sulfur speciation-triggered depletion in ^34^S in vent sourced dietary sources (Fig. 2). Moreover, isotope values of vent crabs in the WV and YV habitats were indistinguishable despite the vast differences in biogeochemical conditions (i.e. sulfur sources) and associated food sources. These multiple lines of evidence show that vent crabs have a confined dietary niche that is much more closely associated with the hydrothermal vent sources than previously recognized.

Compared to the vent crabs, the mollusks at the WV consistently consumed more photo-than chemosynthetic sources. Before the extreme events, the mollusks’ *δ*^13^C values (−20.5‰) were more negative than those of the crabs (−15.3‰). Since seawater POM was −23‰ and vent fluid POM was −17‰ (*11*), our study confirms previously published data that the mollusks relied less on chemosynthetic sources than the vent crabs (*11, 14*). After the extreme events, mollusks gained more access to photosynthetic dietary sources because the resource base at the WV became more influenced by pelagic resources (Fig. 2 and Fig 5). The marked increases in *δ* ^34^S values in all mollusks (average of ∼12.3‰) completely diverged from those of the vent crabs, closely resembling values of pelagic fish and benthic organisms in the open ocean (Fig. 4B and Fig.5). This suggests that mollusks relied more on photosynthetic sources (i.e. seawater sulfate, ∼20‰) than chemosynthetic vent sources (−1‰) (Fig. 5). Indeed, a recent biogeochemical time-series in the region showed a significant decrease in the intensity of conspicuous white color produced by the dominant chemoautotrophic sulfur bacteria (e.g. epsilonproteobacteria and gammaproteobacteria) in the WV seawater after the extreme events (*9*). Decreased consumption of vent associated diet in mollusks is likely due to the lowered contribution of sulfur-based chemotrophic bacteria and vent POM to intermediate consumers such as zooplankton and epi-benthic crustaceans, which make up a large dietary contribution to mollusks (*11*).

We posit that cyanobacteria made greater dietary contributions to the WV food web after the extreme events because of the increasing *δ*^13^C values of the mollusks from 2015 to 2018. We rule out the possibility that increased *δ*^13^C values are directly linked to vent POM sources since the *δ*^15^N values of half of the suspension feeders is close to 0‰, the typical value for biomass from dinitrogen fixation (*34*). Among all marine phytoplankton, the nanoplanktonic cyanobacteria *Trichodesmium*, a diazotroph, has the lowest *δ*^15^N value and highest *δ*^13^C (*34*). In fact, *Trichodesmium* is highly abundant in the Kuroshio Current (*35, 36*), which directly flows through the KST area. Before the extreme events, cyanobacterial production in the KST area was suppressed by the high concentration of toxic metals such as arsenic, cadmium, copper and lead (*37*). The sharp decrease in toxic metals after the extreme events, however, could have led to a surge in cyanobacterial production in the KST vent area (*9*). We attribute the large *δ*^15^N variability (5-6‰) of the suspension feeders to variable contributions of cyanobacterial POM, not the differences in their body sizes because individual specimen sizes of each species were similar (SI Table S1). Interestingly, the two species appear to use similar niches despite their different filtering modes; *T. adamsii* secretes a mucus-like net to filter organic particles in the water, and the free living *B. gravispinosus* filters and forages on organic matter directly from the water column (*20*).

With the caveat that our sample number is small (n = 3), our results indicate intraspecific niche segregation among similar size classes, but further studies are warranted to investigate this hypothesis.

Despite the putative contribution of cyanobacterial production to the KST food web after the disturbance to vent system, both *δ*^13^C and *δ*^15^N values of carnivorous *E. contracta* increased suggesting that they increased their trophic position, probably by feeding on carrion such as zooplankton, vent crab and fish. *E. contracta* likely expanded its foraging area towards the YV area after the decreased toxicity because its *δ*^34^S range widened as opposed to its *δ*^13^C and *δ*^15^N ranges. Taken together, our study shows that each different isotope provided unique insight into changes of macrofauna’s resource utilization in the KST vents. This underpins the importance of using multiple isotope tracers for obtaining a holistic understanding of how drastic biogeochemical changes affects nutrient variability in the hydrothermal vent area.

### Foraging adaptations of endemic and non-endemic species

Vent crabs have until now only been found at three shallow hydrothermal vents with high sulfide concentration (*38*). Hence, vent crabs are considered endemic to these habitats because they are physiologically adapted to tolerate the high concentration of toxic sulfur fluids and are able to ingest sulfur bacteria (*12, 22*). Our results show that vent crabs have an obligate nutritional relationship with chemosynthetic bacteria despite their otherwise opportunistic foraging behavior. The vent crabs’ ability to exploit both chemo- and photosynthetic sources and cope with dramatic disturbances follows their adaptation to an extreme and variable biogeochemical environment over geological timescales. Yet, it appears that the crabs continued to rely more on chemo-than photosynthetic derived sources despite the greatly reduced habitat for chemosynthetic bacteria after YV became less active. The decline in vent crab population with the decrease in the YV activities further underlines that vent crabs are nutritionally dependent on sulfuric bacteria. We argue that this decline is driven by the reduction of chemosynthetic bacteria rather than a higher predation pressure because the vent area still remains toxic and inaccessible to predators after the disturbance. We assume that the crabs feed directly on sulfur bacteria, but it has also been posited that crabs can receive nutritional supplementation from symbiotrophic chemosynthetic bacteria (*23*). This nutritional dependency suggests that vent crabs has low dietary plasticity, and their population will continue to decline if the KST vent activities decrease and become less sulfur-rich.

The three mollusk species we studied in the KST vent area are widely distributed in different world oceans (i.e. Indian or Pacific Oceans). Hence, these non-endemic species are diametrically opposite to the endemic vent crab in terms of their ability to exploit a wide range of resources and inhabit different environments. We show that the food sources of the suspension feeders were strongly affected by the biogeochemical disruption, and that they are less constrained in their exploitation of different chemo- and photosynthetic sources than the vent crabs. After this disruption, the suspension feeders were able to rely on new dietary sources such as cyanobacteria into their diet, and the carnivorous mollusks appeared to increase their dietary breadth by foraging closer to the YV area. The mollusks’ abundance and body sizes did not change after the disturbance show that they are not nutritionally dependent on sulfur bacteria from the vent (*19*). Together with their tolerance to low and medium toxic environments, their wide dietary niche probably plays an important role in their ability to colonize the transitional environment between the extreme hydrothermal vent and open pelagic habitats. Our study furthers our understanding of how these pioneer species are able to inhabit many marine environments in the world’s oceans.

To our knowledge, our study is the first to investigate how sudden and drastic biogeochemical changes affect a unique hydrothermal vent food web. We found that the two suspension feeders exhibited a similar response to the disturbance that is very different from vent crab and carnivorous gastropods. Since the two sampling occasions took place in same season, it is likely that the changes observed in this study truly reflect the impact of extreme events. However, to further understand how vent associated animals respond to extreme events it would be necessary to set up more frequent and longer-term monitoring programs as well as more expansive species sampling in future studies. This could provide more robust inferences in regards to niche segregation among these suspension feeders.

### Foraging by benthic macrofauna under frequent natural and human impact

The diverse response of different macrofauna in our study highlights the importance of understanding ecological heterogeneity and the consequences of habitat disturbance to ecosystem functioning. While the generalist non-endemic species readily adjusted their foraging strategies to the biogeochemical changes in the KST area, the endemic specialist continued to rely on one unique dietary component, sulfur bacteria. This shows that the vent crabs not only occupy habitats inhospitable to other faunal species because it provides them protection against predation, but also that they have formed a nutritional dependency on sulfur bacteria. Similarly close associations with sulfur bacteria have also been found in many deep-sea hydrothermal endemic species such shrimps, vestimentiferan tube worms, bivalves, and gastropods (Fig. 5). The large endemic faunal diversity in deep-sea hydrothermal habitats can undoubtedly be linked to their relatively stable and static environmental conditions. In the much more unstable and dynamic shallow vents, the vent crabs can survive prolonged periods of food scarcity by storing nutrients and exploiting a range of non-sulfur bacterial derived resources (*22*). Nevertheless, the sparse geographical distribution of vent crabs and their population decline following the decrease in YV activities at the KST shows that they are obligate sulfur vent species and that they are much more vulnerable to vent shutdown than the non-endemic mollusks. Our finding is important for assessing the ecological ramifications of human activities, i.e. higher CO_2_ levels, uncontrolled human development, deep-sea mining and industrial dredging, because the alterations of benthic habitats may pass a threshold after which they become uninhabitable for many endemic benthic species that are obligate to unique ecosystems. In such a scenario, generalist species with broad dietary niches and tolerance to rapid changes in biogeochemistry will become more competitive under frequent and severe disturbances from natural hazards and human activities, however, this may come at the cost of substantial loss in biodiversity.

In conclusion, our study demonstrates that extreme events can dramatically disrupt the biogeochemical cycling of shallow marine hydrothermal ecosystems, consequently affecting the relative food basal supplies from photosynthetic and chemosynthetic origins. Our multiple isotopic results show species-specific response to extreme events at two distinct habitats. Non-endemic species show broader dietary flexibility than the endemic species, and thus are more resilient to biogeochemical disruptions. More work must isolate other spatially fragmented habitats containing macrofauna assemblages comprising rare, endangered, or endemic species. Specifically, long-term observations dedicated to monitoring marine habitats that have experienced extraordinary amounts of human use and exploitation will become increasingly necessary to track ecosystem health. Our work exemplifies how marine specialists are more vulnerable than generalists to the unprecedented evolutionary pressure exerted by human activities worldwide.

## Materials and Methods

### Sampling and processing

The research cruises for the collection of vent organisms were carried out between April and May each year in 2015 and 2018 to avoid seasonal effects on food source and seawater biogeochemistry, which is strongly influenced by sea water temperature and the direction of the predominant tidal and Kuroshio Current. No field sampling campaign took place in May 2016 due to the natural disasters. After the extreme events, the YV landscape was fragmented due to the YV’s close proximity to the shore of KST. The accumulated YV sulfur chimney was completely destroyed and the original seabed landscape was buried with new layers of debris and sediments. The vent activities in YV increased in strength again in 2018 in many smaller fumaroles. Therefore, the subsequent sampling in 2018 for YV was conducted on top of the buried old YV site, located through the GPS and other landmarks. The WV seabed, which is farther away from the shoreline than the YV area, was only buried with a thin layer of sediment and exhibited less venting activity in 2018 than 2015. This situation enabled us to sample the same site in the YV in 2018.

Vent organisms were collected on each cruise to the YV and WV via SCUBA diving by a minimum of two scientific divers. We did not collect organisms outside of the YV and WV areas since they remained mostly unaffected by the vent chemistry in the mixing zone, and the seafloor fauna typically reflect the surrounding shelf benthic habitats (*19*). Vent crabs (*X testudinatus*) were collected at both the YV and WV. In 2018, collected crabs were sexed based on their shape of pleonal structures and males and females were processed separately. Key species of mollusks including sessile (*T. adamsii*) and motile gastropod (*B. gravispinosu* and *E. contracta*) were collected at the WV within a 10 m radius within the same site each year. For statistical reasons, at least 6 specimens of each species were collected. All organisms were placed in portable nets and brought to the surface where they were then placed in aerated containers with seawater. After sampling, vent invertebrates were transported to the laboratory at National Taiwan Ocean University, and placed in an 80 × 50 × 60 (L×W× H) tank with a constant fresh filtered seawater flow for 24 h to empty their gut contents. However, the filter feeders, namely, *T. adamsii* and *B. gravispinosu*, died or were preyed on by carnivorous *E. contracta* during the transport and 24 h gut cleaning. Dead specimens were discarded and resulted in a smaller sample size of the filter feeders. In 2015, due to lack of trained personnel on-site for dissecting the animals, all vent invertebrates were rinsed in de-ionized water and then were frozen at −20°C before drying in the oven at 60°C for 48 h. The dry samples were then shipped back and dissected in the Institute of Geosciences, Kiel University (CAU). In 2018, after being rinsed in de-ionized water, all molluscan samples were frozen at −20°C for euthanization and the crabs were desensitized by placing them in ice for half hour before dissection. For the crabs, we pooled and analyzed muscle tissues from the claws. For the mollusks, we used all muscle tissue available including foot and abductor muscles due to the small mass of the specimens. To investigate whether the macrofanua species changed their body sizes in response to the biogeochemical changes, we also measured the sizes of vent crabs and mollusks except for the irregularly coiled *T. adamsii*. We determined the size of vent crab by measuring the distance between the two most protruding notches. We used the shell height and length, respectively, to determine the body sizes of the gastropod *E. contracta* and *B. gravispinosu* was determined by. All the body size data is presented in the supplementary Table S1. At the Institute of Geosciences in CAU, all samples were homogenized using a mortar and pestle and subdivided into two fractions, one for C and N stable isotope analyses, and the other for S stable isotope analyses.

### Stable carbon and nitrogen isotope analyses

For the samples collected in 2015, approximately 60 to 100 µg of dry mass for each sample were weighed into tin capsules. Samples were then analyzed for stable C and N isotope ratios using a continuous-flow isotope-ratio mass spectrometer MAT253 (Thermo Finnigan) coupled to an elemental analyzer EA1108 (Carlo Erba Instruments) (EA-IRMS) through a Conflo III interface (ThermoFinnigan) in the stable isotope facility of the University of La Coruña, Spain. During the analyses, a set of international reference materials were used for isotope calibration (NBS 22, IAEA-CH-6, USGS24 for *δ*^13^C and IAEA-N-1, IAEA-N-2, IAEA-NO-3 for *δ*^15^ N). The analytical measurement errors of ± 0.15 ‰ were calculated for *δ*^13^C and *δ*^15^N based on the replicated assays of the laboratory standard acetanilide interspersed between sample analyses. Samples collected in 2018 were measured with an EA-IRMS in the Iso Analytical Limited Inc, UK. Approximately 1 milligram of the dry mass of each sample was used for the carbon and nitrogen isotope analysis. For quality control, internal lab standards (IA-R068, IA-R038, IA-R069, and a mixture of IAEA-C7 and IA-R-R046) were analyzed in between sample runs. These standards were calibrated against international reference material IAEA-CH-6, IAEA-N-1, IAEA-C-7 for both *δ*^13^C and *δ*^15^N. The isotope data are also expressed in delta (*δ*) *permil* (‰): *δ* (‰) = [(*R*_sample_/*R*_standard_) – 1] ×1000, where *R* is the ratio of heavy to light isotope. The isotope ratios are expressed relative to international standards: VPDB for carbon and atmospheric air for nitrogen. For organisms C: N ratios greater than 3.5 in this study, we applied lipid correction to *δ*^13^C values for muscle tissue samples (*N*=49) following Logan et al. (*39*). We did not perform lipid extraction prior to stable isotope analyses of muscle tissues because lipid extraction is known to affect *δ*^15^N values (*40*).

### Stable sulfur isotope analyses

Sulfur isotope analyses of all samples were performed on an EA-IRMS at Iso-Analytical Limited, UK. Tin capsules containing reference or sample material plus vanadium pentoxide catalyst were loaded into an automatic sampler and dropped, in sequence, into a furnace held at 1080 °C, and combusted with oxygen. For quality control purposes, test samples of a set of in-house standards IA-R061 (barium sulfate) (N=12), IAEA-SO-5 (barium sulfate) (N=5), IA-R068 (soy protein) (N=5) and IA-R069 (tuna protein) (N=3) were measured during batch analysis of our samples.

Internal laboratory standards, IA-R061, IA-R025 (barium sulfate) and IA-R026 (silver sulfide) were used for calibration and correction of ^18^O contribution to the SO^+^ ion beam. Calibration of internal standards were carried out against inter-laboratory comparison standards (NBS-127, barium sulfate and IAEA-S-1, silver sulfide) distributed by the International Atomic Energy Agency. The standard deviation for the test samples were ± 0.10‰ except for IA-R069 (±0.32‰). The sulfur isotope ratios also expressed in delta (*δ*) *permil* (‰): *δ* (‰) = [(*R*_sample_/*R*_standard_) – 1] ×1000 and relative to international standards: Vienna Canyon Diablo Troilite (V-CTD) for sulfur.

### Vent crab population density data change from 2004 to 2018

We examined the relationship between vent activity and crab abundance to understand whether the decreased vent activity affected the vent crab population. We conducted the underwater video survey next to the new Yellow Vent mouths, the most populous area for vent crabs. A digital camera was set up to capture the 50×88 cm^2^ area for three days, from August 10^th^ to August 12^th^, 2018. To be consistent with previous published population data, which was only captured during the day time, we only used the footage from the daylight hours for population density analysis. In addition, our study area is subjected to strong tidal currents mixing with vent fluid discharge, which make the water too murky for clear visualization after 15:00. Therefore, we used images captured between 10:00 to 15:00. We omitted film footage from 11:00 onwards on August 12^th^ because the image quality was blurry, possibly due to a technical failure of the camera equipment or strong mixing of the underwater currents. To count for the mobile individuals which entered the survey area, we split each hour into six 10 minute intervals, which amounted to 66 film frames for the all the hours that were used for this analysis. Then we compared with the vent crab population baseline established in 2004 and 2014 (*13, 19*). To make the vent crab density data comparable for three years, we have converted our own count and published data to area coverage of 25×25 cm^2^ based on Chan et al. (2016). In 2004, the average density count was 364 specimens for every 100 cm × 100 cm at the YV, which is the equivalent of 23 specimens for every 25×25 cm^2^ (*13*). By 2014, the average speciemens was 11 specimens at the YV for every 25cm × 25cm quadrats (*19*).

### Statistical analyses

All statistical analyses were performed in R version 3.3.4 (R-Development-Core-Team, 2017-11-30) with R-Studio interface version 1.0.136. All isotope data (*δ*^13^C, *δ*^15^N, and *δ*^34^S) were assessed for homoskedasticity (equal-variance among organism groups) by using Fligner-Killeen tests and were visually checked for any departures from normality on the Q-Q plot. We found *P*-values of 0.87, 0.23, 0.29 for *δ*^13^C, *δ*^15^N, and *δ*^34^S, respectively, indicating that the variation in all organism was homogenous. All the isotope values were also assessed (R function: cor) for their correlation with each other, and we found that none of the parameters co-varied with one another. Based on these tests, we carried out statistical analyses on the isotope results. We first tested for significant differences in inter-annual isotope values (*δ*^13^C, *δ*^15^N, and *δ*^34^S) among vent crabs from 2015 and 2018 by applying Pillai’s trace Multivariate Analysis of Variance (MANOVA). To assess which dependent variables (*δ*^13^C, *δ*^34^S and/or *δ*^15^N values) are significantly different between groups (R function: summary.aov), we then used univariate Analysis of Variance (ANOVA) on the output from MANOVA. Instead of using multiple t-tests, we used a MANOVA test to compare the vent crabs from 2015 and 2018, because the assumptions for MANOVA test are satisfied. For the mollusk species, we performed two sample t-tests on isotopic changes on *E. contracta* before and after the extreme events because both the standard population effect size (D>4) and sample sizes satisfied the statistical assumption (*41*). However, no statistical analyses were performed on *B. gravispinosus* and *T. adamsii* due to the small sample numbers (N=3) and their effect size (*41*).

Given that vent crabs are considered an endemic species in the KST, we examined whether or not there was a significant difference in the intra-annual variability of isotope values (*δ*^13^C, *δ*^15^N, and *δ*^34^S) between vent crabs and the pooled mollusk species by performing MANOVA and ANOVA analyses. We compared all three isotope variables (*δ*^13^C, *δ*^15^N, and *δ*^34^S) for 2015 but only *δ*^13^C and *δ*^34^S for 2018 as *δ*^15^N value ranges of mollusks in 2018 were wider compared to those of vent crabs. MANOVA and ANOVA tests were also conducted to examine whether there were any biases in the vent crab’s isotope values between habitats (i.e. YV vs. WV, N=6 and 10, respectively), as well as between the different sexes (female vs. male, N=8 for each) in 2018. Finally, we compared the body size changes before and after the extreme events using two sample t-test. The *P*-adjusted values from all ANOVA decomposition tests were obtained using *p*.*adjust* function with FRD method.

Unless otherwise stated, statistical significance is assessed at *P*< 0.05. All figures are made using “ggplot2” and “ggpubr” packages.

### Comparison of δ^34^S values between hydrothermal vent organisms and marine organisms in the open ocean

To gain insight how *δ*^34^S values of the KST vent macrofauna compared to those deep-sea hydrothermal vent organisms as well as marine organisms in the open ocean, we compiled and compared the *δ*^34^S data from this study with *δ*^34^S data from the previously published literature (Fig. 5): for example, jellyfish from the UK shelf (*42*); pelagic fish (*43*); offshore_benthic_invertebrates (*43-46*), and as well as deep sea hydrothermal vent animals (*45, 47, 48*). We also compared *δ*^34^S values of the KST vent macrofauna with that of KST native sulfur and seawater sulfate (*21*). This will provide a larger context for understanding and interpreting the *δ*^34^S results presented in this study.

## Data Availability

The data that support the findings of this study are provided in Supplementary table S1.

## Acknowledgements

We thank caption Rongji Zhang for his support for our cruise and Shih-hao Huang for field assistance. We are grateful to Michael J. Storozum for editorial comments and valuable discussion that improved the manuscript.

## Author Contributions

Y.V.W., M. L., D.G-S., L-C. and J-S. H. designed the work. Y. V. W., M. L., L-C. T., P-W. L, and N. S-S conducted the field work and collected samples. Y. V. W, N. S. S, and P-W. L conducted the lab work. Y. V. W. and T. L. analyzed the isotope data. T.Y. C. provided the underwater digital film recording in 2018; P-W. L analyzed the film data and calculated the vent crab numbers. Y.V.W. and T. L. wrote the manuscript. All authors commented on the manuscripts. Cruises were coordinated by: M.L., D.G-S., L-C.T., and J-S.H.

## Data and material availability

All the raw data from this study is included in the Supplementary Materials. Arial photo images associated with this study is available under Accession Number 0175781 in https://data.nodc.noaa.gov/cgi-bin/iso?id=gov.noaa.nodc:0175781 with DOI: 10.25921/6hy3-6d56.

## Additional Information

Supplementary information is available in the online version of the paper.

## Supplementary Materials

### Supplementary text 1: Background on shallow marine vent food sources

Together with deep sea vents, shallow vents are unique underwater volcanic features that, despite their extreme fluid chemical properties, facilitate habitat formation by creating new niches for vent specialists and pioneer species in otherwise homogenous and poor habitats (*8, 19, 20, 28*). These shallow vents are home to a diverse community of bentho-pelagic species that are primarily supported by chemical energy created when warm vent fluids rich in sulfides, methane, and other minerals meet seawater (*4, 8, 15, 19, 20*). In addition to the chemosynthetic sources, hydrothermal benthic and planktonic heterotrophic communities are also adapted to use dietary sources of photosynthetic origin. The extent to which these two dietary resources are available largely depends on vent intensity and its interaction with surrounding ecosystems through horizontal and vertical transport pathways that are affected by multifaceted factors (*15*). Some of these factors such as fluid flow, seawater mixing and entraining, biogeochemical process, and biological succession are highly sensitive to perturbations (*8, 15, 18*).

### Supplementary text II: Background of the study region

Kueishantao Island (KST, 24.843°N, 121.951°E), also known as Turtle Island, is situated at a tectonic junction off the coast of northeastern Taiwan and the southern end of the Okinawa Trough. KST is a Holocene stratovolcano and volcanic activity beneath the KST area is still vigorous, even though the last eruption occurred ∼7000 years ago (*21*). In the last two decades, KST shallow vents are among the most intensively studied shallow vent systems in the world due to their extreme geochemical properties and their easy accessibility. The active YV vents are typically within the temperature range of >70 °C and fluid fluxes are up to 150 m^3^/h, whereas the temperature range of the semi-inactive WV fluctuates between 30– 65 °C, with frequent degassing activity (*21, 27*). The low pH values (down to 1.75 at certain locations) were recorded as the lowest in the world two decades ago (*21*). Before May 2016, more than 30 hydrothermal vents located at a water depth between 6 to 30 m were still active (*21, 27*), making them easily accessible for scientists to study the geochemical and biological processes of these vent systems. However, in 2016, KST was hit by a M5.8 earthquake and a subsequent C5 typhoon within a few weeks (12^th^ May and 2^nd^-10^th^ July, respectively), disrupting normal conditions, and providing a unique opportunity to study food web changes induced by the extreme events.

---

**Supplementary Table S1.** Individual *δ*^13^C, *δ*^15^N and *δ*^34^S values of individual organisms and the shell width from the Kueishantao (KST) shallow hydrothermal vent.

| Species | Body size(mm) | Year | $\delta^{13}\text{C}$ | $\delta^{15}\text{N}$ | $\delta^{34}\text{S}$ |
| --- | --- | --- | --- | --- | --- |
| *Bostrycapulus gravispinosus* | 14.24 | 2015 | -20.2 | 6.9 | 10.3 |
| *Bostrycapulus gravispinosus* | 13.92 | 2015 | -20.4 | 6.8 | 8.2 |
| *Bostrycapulus gravispinosus* | 13.77 | 2015 | -20.4 | 6.4 | 8.7 |
| *Ergalatax contracta* | 24.12 | 2015 | -19.2 | 8.2 | 7.3 |
| *Ergalatax contracta* | 22.62 | 2015 | -19.7 | 8.6 | 6.4 |
| *Ergalatax contracta* | 20.20 | 2015 | -20.3 | 7.8 | 6.7 |
| *Thylacodes adamsii* | N.D | 2015 | -21.4 | 7.4 | 4.8 |
| *Thylacodes adamsii* | N.D | 2015 | -21.6 | 7.1 | 7.5 |
| *Thylacodes adamsii* | N.D | 2015 | -21.1 | 6.6 | 9.4 |
| *Xenograpsus testudinatus* | 23.58 | 2015 | -13.1 | 5.0 | 4.3 |
| *Xenograpsus testudinatus* | 22.89 | 2015 | -11.1 | 5.2 | 6.9 |
| *Xenograpsus testudinatus* | 21.12 | 2015 | -13.6 | 5.3 | 6.3 |
| *Xenograpsus testudinatus* | 18.36 | 2015 | -16.8 | 6.1 | 10.0 |
| *Xenograpsus testudinatus* | 21.91 | 2015 | -16.9 | 6.7 | 10.3 |
| *Xenograpsus testudinatus* | 22.99 | 2015 | -15.6 | 5.2 | 9.4 |
| *Xenograpsus testudinatus* | 23.18 | 2015 | -15.5 | 4.0 | 7.7 |
| *Xenograpsus testudinatus* | 19.46 | 2015 | -15.7 | 5.7 | 10.3 |
| *Xenograpsus testudinatus* | 26.86 | 2015 | -18.1 | 5.3 | 9.1 |
| *Xenograpsus testudinatus* | 19.54 | 2015 | -18.0 | 6.7 | 7.9 |
| *Xenograpsus testudinatus* | 20.89 | 2015 | -16.1 | 4.3 | 3.9 |
| *Xenograpsus testudinatus* | 16.36 | 2015 | -13.2 | 5.2 | 7.6 |
| *Xenograpsus testudinatus* | 22.83 | 2018 | -15.7 | 8.6 | 5.8 |
| *Xenograpsus testudinatus* | 18.84 | 2018 | -16.9 | 7.8 | 6.7 |
| *Xenograpsus testudinatus* | 23.84 | 2018 | -18.2 | 8.6 | 5.8 |
| *Xenograpsus testudinatus* | 22.46 | 2018 | -15.3 | 8.2 | 5.3 |
| *Xenograpsus testudinatus* | 18.69 | 2018 | -17.6 | 7.5 | 4.7 |
| *Xenograpsus testudinatus* | 15.96 | 2018 | -16.3 | 6.0 | 1.2 |
| *Xenograpsus testudinatus* | 24.31 | 2018 | -17.2 | 8.2 | 4.0 |
| *Xenograpsus testudinatus* | 18.09 | 2018 | -16.5 | 7.8 | 8.0 |
| *Xenograpsus testudinatus* | 22.18 | 2018 | -15.6 | 6.4 | 4.8 |
| *Xenograpsus testudinatus* | 18.17 | 2018 | -17.8 | 8.1 | 5.7 |
| *Xenograpsus testudinatus* | 27.14 | 2018 | -13.8 | 6.7 | 5.3 |
| *Xenograpsus testudinatus* | 18.73 | 2018 | -15.7 | 7.1 | 5.9 |
| *Xenograpsus testudinatus* | 19.16 | 2018 | -15.2 | 7.1 | 7.0 |
| *Xenograpsus testudinatus* | 17.32 | 2018 | -15.5 | 6.9 | 5.7 |
| *Xenograpsus testudinatus* | 19.53 | 2018 | -15.9 | 7.4 | 5.4 |
| *Xenograpsus testudinatus* | 18.85 | 2018 | -15.5 | 7.4 | 5.8 |
| *Ergalatax contracta* | 20.85 | 2018 | -15.3 | 9.1 | 7.9 |
| *Ergalatax contracta* | 22.8 | 2018 | -15.2 | 10.5 | 9.5 |
| *Ergalatax contracta* | 18.0 | 2018 | -15.5 | 9.6 | 9.0 |
| *Ergalatax contracta* | 21.47 | 2018 | -16.2 | 10.2 | 12.6 |

| **Species** | **Body size(mm)** | **Year** | **$\delta^{13}\text{C}$** | **$\delta^{15}\text{N}$** | **$\delta^{34}\text{S}$** |
| --- | --- | --- | --- | --- | --- |
| *Ergalatax contracta* | 20.44 | 2018 | -16.0 | 9.8 |  |
| *Ergalatax contracta* | 20.0 | 2018 | -15.2 | 10.2 | 10.7 |
| *Thylacodes adamsii* | N.D | 2018 | -18.7 | 5.9 | 11.0 |
| *Thylacodes adamsii* | N.D | 2018 | -18.2 | 0.7 | 13.8 |
| *Thylacodes adamsii* | N.D | 2018 | -18.6 | 5.7 | 13.8 |
| *Bostrycapulus gravispinosus* | 12.98 | 2018 | -18.2 | 4.6 | 11.7 |
| *Bostrycapulus gravispinosus* | 13.34 | 2018 | -16.6 | 0.3 | 12.4 |
| *Bostrycapulus gravispinosus* | 13.35 | 2018 | -17.6 | 1.2 | 11.5 |

**Supplementary Table S2.** MANOVA and ANOVA results comparing *δ*^13^C, *δ*^15^N, *δ*^34^S values between 2015 and 2018 for vent crab (*Xenograpsus testudinatus*).

Manova summary
|  | Df | Pillai | approx. F | num Df | den Df | P-adj |
| --- | --- | --- | --- | --- | --- | --- |
| Regression | 1 | 0.889 | 50.8 | 3 | 19 | 6.397e-09 *** |
| Residuals | 21 |  |  |  |  |  |

ANOVA summary Response $\delta^{34}\text{S}$ :
|  | Df | Sum Sq | Mean Sq | F value | P-adj |
| --- | --- | --- | --- | --- | --- |
| Sampling_year | 1 | 70.283 | 70.283 | 37.464 | 0.003 ** |
| Residuals | 21 | 39.397 | 1.876 |  |  |

Response $\delta^{13}\text{C}$ :
|  | Df | Sum Sq | Mean Sq | F value | P-adj |
| --- | --- | --- | --- | --- | --- |
| Sampling_year | 1 | 1.1615 | 1.1615 | 0.906 | 0.1822 |
| Residuals | 21 | 26.9315 | 1.2825 |  |  |

Response $\delta^{15}\text{N}$ :
|  | Df | Sum Sq | Mean Sq | F value | P-adj |
| --- | --- | --- | --- | --- | --- |
| Sampling_year | 1 | 16.060 | 16.0604 | 23.765 | 6.51e-07 *** |
| Residuals | 21 | 14.192 | 0.6758 |  |  |

**Supplementary Table S3.** Summary of Welch Two Sample t-test comparing *δ*^13^C, *δ*^15^N, and *δ*^34^S values for *Ergalatax contracta* before and after disturbance. Lower and upper limits of 95% Confidence Intervals (CI) are also presented. The isotope value unit is per mil (‰).

|  | Mean of 2018 | Mean of 2015 | CI.lower | CI.upper | p-adj value |
| --- | --- | --- | --- | --- | --- |
| $\delta^{34}\text{S}$ | 9.94 | 6.8 | 0.94 | 5.34 | 0.015 |
| $\delta^{13}\text{C}$ | -19.73 | -15.57 | 3.07 | 5.27 | 0.003 |
| $\delta^{15}\text{N}$ | 9.90 | 8.20 | 0.91 | 2.49 | 0.004 |

**Supplementary Table S4.**
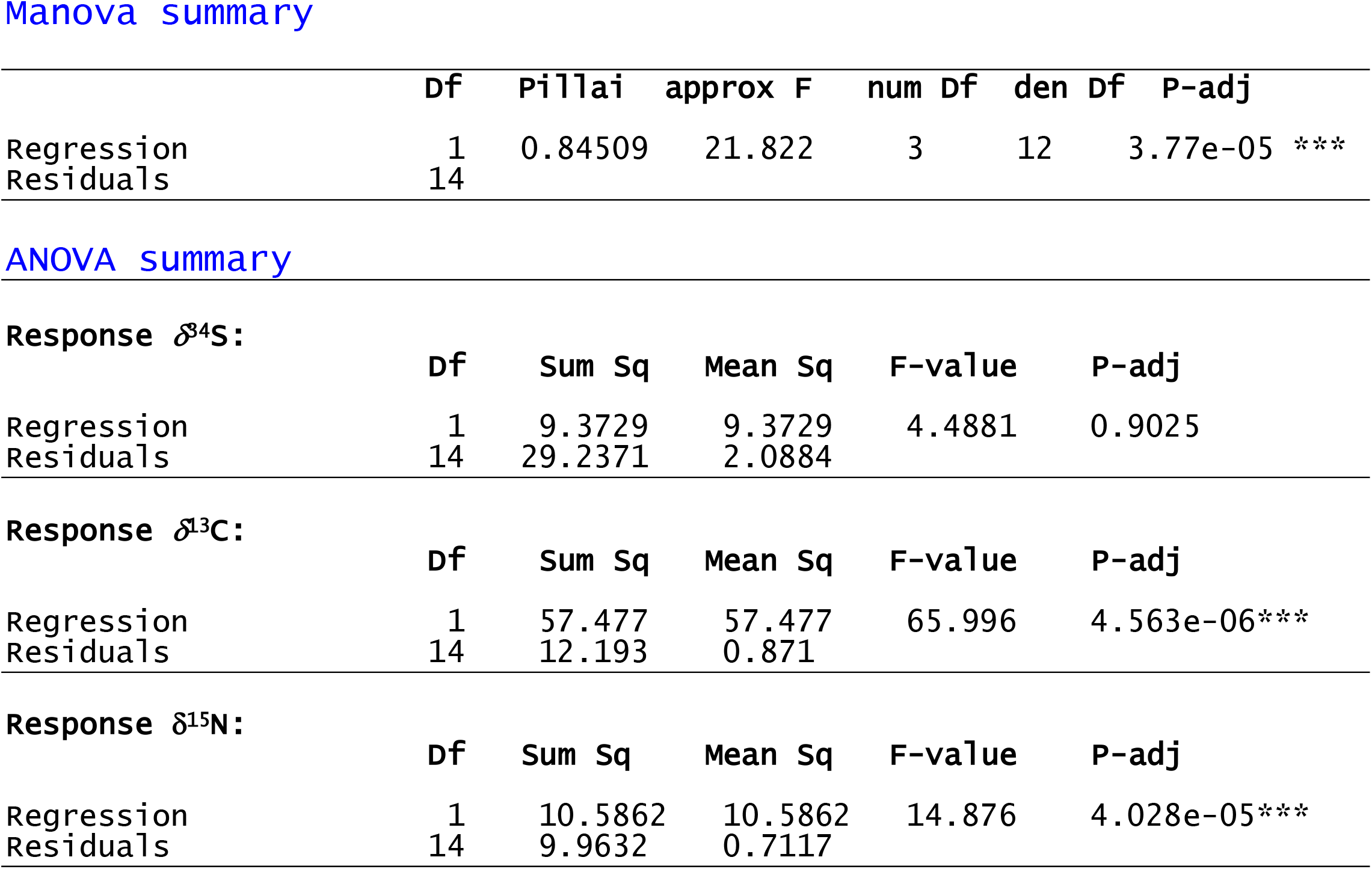
MANOVA and ANOVA results comparing *δ*^13^C, *δ*^15^N, *δ*^34^S values between the vent crab (*Xenograpsus testudinatus*) and pooled Mollusca group (*Ergalatax contracta, Thylacodes adamsii, Bostrycapulus gravispinosus*) in 2015.

**Supplementary Table S5.** MANOVA and ANOVA results comparing *δ*^13^C, *δ*^34^S values between the vent crab (*Xenograpsus testudinatus*) and pooled Mollusca group (*Ergalatax contracta, Thylacodes adamsii, Bostrycapulus gravispinosus*) in 2018.

Manova summary
|  | Df | Pillai | approx F | num Df | den Df | P-adj |
| --- | --- | --- | --- | --- | --- | --- |
| Regression | 1 | 0.76966 | 40.096 | 2 | 24 | 2.231e-08 *** |
| Residuals | 25 |  |  |  |  |  |

ANOVA summary
| Response $\delta^{34}\text{S}$: | | | | | |
| --- | --- | --- | --- | --- | --- |
|  | Df | Sum Sq | Mean Sq | F-value | P-adj |
| Regression | 1 | 220.789 | 220.789 | 80.598 | 5.396e-09*** |
| Residuals | 25 | 68.485 | 2.739 |  |  |

| Response $\delta^{13}\text{C}$: | | | | | |
| --- | --- | --- | --- | --- | --- |
|  | Df | Sum Sq | Mean Sq | F-value | P-adj |
| Regression | 1 | 2.985 | 2.9850 | 1.8453 | 0.1865 |
| Residuals | 25 | 40.442 | 1.6177 |  |  |

**Supplementary Table S6.** MANOVA and ANOVA results comparing *δ*^13^C, *δ*^15^N, and *δ*^34^S values between the *Xenograpsus testudinatus* collected in the Yellow and White Vents in 2018 (N=16). This result shows that there are no significant differences in *δ*^13^C, *δ*^15^N, and *δ*^34^S for vent crabs in different habitats.

Manova summary
|  | Df | Pillai | approx F | num Df | den Df | P-adj |
| --- | --- | --- | --- | --- | --- | --- |
| Regression | 1 | 0.48737 | 3.803 | 3 | 12 | 0.0796 |
| Residuals | 14 |  |  |  |  |  |

ANOVA summary
| Response $\delta^{34}\text{S}$: | | | | | |
| --- | --- | --- | --- | --- | --- |
|  | Df | Sum Sq | Mean Sq | F value | P-adj |
| Location | 1 | 1.5844 | 1.584 | 0.7208 | 0.4102 |
| Residuals | 14 | 30.775 | 2.198 |  |  |

| Response $\delta^{13}\text{C}$: | | | | | |
| --- | --- | --- | --- | --- | --- |
|  | Df | Sum Sq | Mean Sq | F value | P-adj |
| Location | 1 | 7.812 | 7.812 | 9.282 | 0.0348 |
| Residuals | 14 | 11.782 | 0.8416 |  |  |

| Response $\delta^{15}\text{N}$: | | | | | |
| --- | --- | --- | --- | --- | --- |
|  | Df | Sum Sq | Mean Sq | F value | P-adj |
| Location | 1 | 1.442 | 1.442 | 2.766 | 0.1580 |
| Residuals | 14 | 7.296 | 0.521 |  |  |

**Supplementary Table S7.** MANOVA and ANOVA results comparing *δ*^13^C, *δ*^15^N, and *δ*^34^S values between the male and female *Xenograpsus testudinatus* in 2018 (N=16). The results show that there is no significant difference in *δ*^13^C, *δ*^15^N, and *δ*^34^S values between sexes.

|  | Df | Pillai | approx F | num Df | den Df | P-adj |
| --- | --- | --- | --- | --- | --- | --- |
| Regression | 1 | 0.2716 | 1.4915 | 3 | 12 | 0.2667 |
| Residuals | 14 |  |  |  |  |  |

ANOVA summary
| Response $\delta^{34}\text{S}$: | | | | | |
| --- | --- | --- | --- | --- | --- |
|  | Df | Sum Sq | Mean Sq | F value | P-adj |
| Sex | 1 | 0.031 | 0.0306 | 0.0133 | 0.91 |
| Residuals | 14 | 32.329 | 2.3092 |  |  |

| Response $\delta^{13}\text{C}$: | | | | | |
| --- | --- | --- | --- | --- | --- |
|  | Df | Sum Sq | Mean Sq | F value | P-adj |
| Sex | 1 | 1.756 | 1.756 | 1.378 | 0.2601 |
| Residuals | 14 | 17.839 | 1.274 |  |  |

| Response $\delta^{15}\text{N}$: | | | | | |
| --- | --- | --- | --- | --- | --- |
|  | Df | Sum Sq | Mean Sq | F value | P-adj |
| Sex | 1 | 0.250 | 0.250 | 0.412 | 0.5311 |
| Residuals | 14 | 8.488 | 0.606 |  |  |

**Supplementary Table S8.** Vent crab population density data in YV from 2004 to 2018.

| **Year** | **Density**<br>**(25×25cm<sup>2</sup>)** | **Reference** |
| --- | --- | --- |
| 2004 | 23 | Jeng et al., 2004 |
| 2014 | 11 | Chan et al., 2016 |
| 2018 | 4 | This study |

## References

1. A. P. Rees, Pressures on the marine environment and the changing climate of ocean biogeochemistry. Philosophical Transactions of the Royal Society A: Mathematical, Physical and Engineering Sciences 370, 5613–5635 (2012).

2. O. S. Ashford et al., On the Influence of Vulnerable Marine Ecosystem Habitats on Peracarid Crustacean Assemblages in the Northwest Atlantic Fisheries Organisation Regulatory Area. Frontiers in Marine Science 6, (2019).

3. M. Gianni et al., How Much Longer Will it Take? A Ten-Year Review of the Implementation of United Nations General Assembly resolutions 61/105, 64/72 and 66/68 on the Management of Bottom Fisheries in Areas Beyond National Jurisdiction: Deep Sea Conservation Coalition., (2016).

4. J. M. Hall-Spencer et al., Volcanic carbon dioxide vents show ecosystem effects of ocean acidification. Nature 454, 96 (2008).

5. Y. Harada et al., Stable isotopes indicate ecosystem restructuring following climate-driven mangrove dieback. Limnology and Oceanography n/a, (2019).

6. M. E. Ledger et al., in Advances in Ecological Research, G. Woodward, E. J. O’Gorman, Eds. (Academic Press, 2013), vol. 48, pp. 343–395.

7. R. Sagarin et al., Between control and complexity: opportunities and challenges for marine mesocosms. Frontiers in Ecology and the Environment 14, 389–396 (2016).

8. V. G. Tarasov, A. V. Gebruk, A. N. Mironov, L. I. Moskalev, Deep-sea and shallow-water hydrothermal vent communities: Two different phenomena? Chem Geol 224, 5–39 (2005).

9. M. Lebrato et al., Earthquake and typhoon trigger unprecedented transient shifts in shallow hydrothermal vents biogeochemistry. Scientific Reports 9, 16926 (2019).

10. H.-U. Dahms, L.-C. Tseng, J.-S. Hwang, Are vent crab behavioral preferences adaptations for habitat choice? PLOS ONE 12, e0182649 (2017).

11. N.-N. Chang et al., Trophic structure and energy flow in a shallow-water hydrothermal vent: Insights from a stable isotope approach. PLOS ONE 13, e0204753 (2018).

12. T. W. Ho, J.-S. Hwang, M. K. Cheung, H. S. Kwan, C. K. Wong, Dietary analysis on the shallow-water hydrothermal vent crab Xenograpsus testudinatus using Illumina sequencing. Mar Biol 162, 1787–1798 (2015).

13. M. S. Jeng, N. K. Ng, P. K. Ng, Feeding behaviour: hydrothermal vent crabs feast on sea ‘snow’. Nature 432, 969 (2004).

14. T.-W. Wang, T.-Y. Chan, B. K. K. Chan, Trophic relationships of hydrothermal vent and non-vent communities in the upper sublittoral and upper bathyal zones off Kueishan Island, Taiwan: a combined morphological, gut content analysis and stable isotope approach. Mar Biol 161, 2447–2463 (2014).

15. L. A. Levin et al., Hydrothermal Vents and Methane Seeps: Rethinking the Sphere of Influence. Frontiers in Marine Science 3, (2016).

16. Z. Minic, Organisms of deep sea hydrothermal vents as a source for studying adaptation and evolution. Symbiosis 47, 121–132 (2009).

17. C. T. S. Little, R. C. Vrijenhoek, Are hydrothermal vent animals living fossils? Trends Ecol Evol 18, 582–588 (2003).

18. E. Ramirez-Llodra et al., Man and the Last Great Wilderness: Human Impact on the Deep Sea. PLOS ONE 6, e22588 (2011).

19. B. K. K. Chan et al., Community Structure of Macrobiota and Environmental Parameters in Shallow Water Hydrothermal Vents off Kueishan Island, Taiwan. PLOS ONE 11, e0148675 (2016).

20. C. Chen, T.-Y. Chan, B. K. K. Chan, Molluscan diversity in shallow water hydrothermal vents off Kueishan Island, Taiwan. Marine Biodiversity, (2017).

21. C.-T. A. Chen et al., Tide-influenced acidic hydrothermal system offshore NE Taiwan. Chem Geol 224, 69–81 (2005).

22. M. Y.-A. Hu, W. Hagen, M.-S. Jeng, R. Saborowski, Metabolic energy demand and food utilization of the hydrothermal vent crab Xenograpsus testudinatus (Crustacea: Brachyura). Aquat Biol 15, 11–25 (2012).

23. S.-H. Yang, P.-W. Chiang, T.-C. Hsu, S.-J. Kao, S.-L. Tang, Bacterial Community Associated with Organs of Shallow Hydrothermal Vent Crab Xenograpsus testudinatus near Kuishan Island, Taiwan. PLOS ONE 11, e0150597 (2016).

24. B. Fry, H. Gest, J. M. Hayes, Sulphur isotopic compositions of deep-sea hydrothermal vent animals. Nature 306, 51–52 (1983).

25. L. G. Woodruff, W. C. Shanks, Sulfur isotope study of chimney minerals and vent fluids from 21°N, East Pacific Rise: Hydrothermal sulfur sources and disequilibrium sulfate reduction. Journal of Geophysical Research: Solid Earth 93, 4562–4572 (1988).

26. C. L. Van Dover, B. Fry, Stable isotopic compositions of hydrothermal vent organisms. Mar Biol 102, 257–263 (1989).

27. S. Oana, H. Ishikawa, Sulfur isotopic fractionation between sulfur and sulfuric acid in the hydrothermal solution of sulfur dioxide. Geochem J 1, 45–50 (1966).

28. A. Ueda, H. Sakai, A. Sasaki, Isotopic composition of volcanic native sulfur from Japan. Geochem J 13, 269–275 (1979).

29. C. E. de Ronde et al., Intra-oceanic subduction-related hydrothermal venting, Kermadec volcanic arc, New Zealand. Earth and Planetary Science Letters 193, 359–369 (2001).

30. H. Sakai, Isotopic properties of sulfur compounds in hydrothermal processes. Geochem J 2, 29–49 (1968).

31. S. V. Galkin, A. M. Sagalevich, in Extreme biomimetics. (Springer, 2017), pp. 97–118.

32. X. Wu, B. Wu, J. Song, X. Li, Spatio-temporal distribution of dissolved sulfide in China marginal seas. Chinese Journal of Oceanology and Limnology 32, 1145–1156 (2014).

33. B. Fry et al., Stable isotope studies of the carbon, nitrogen and sulfur cycles in the Black Sea and the Cariaco Trench. Deep Sea Research Part A. Oceanographic Research Papers 38, S1003–S1019 (1991).

34. E. J. Carpenter, H. R. Harvey, B. Fry, D. G. Capone, Biogeochemical tracers of the marine cyanobacterium Trichodesmium. Deep Sea Research Part I: Oceanographic Research Papers 44, 27–38 (1997).

35. T. Shiozaki et al., Why is Trichodesmium abundant in the Kuroshio? Biogeosciences 12, 6931–6943 (2015).

36. C. Wu, F.-X. Fu, J. Sun, S. Thangaraj, L. Pujari, Nitrogen Fixation by Trichodesmium and unicellular diazotrophs in the northern South China Sea and the Kuroshio in summer. Scientific Reports 8, 2415 (2018).

37. S. Yadav, R. Prajapati, N. Atri, Effects of UV-B and heavy metals on nitrogen and phosphorus metabolism in three cyanobacteria. Journal of basic microbiology 56, 2–13 (2016).

38. H. Miyake, A. Oda, S. Wada, T. Kodaka, S. Kurosawa, First record of a shallow hydrothermal vent crab, Xenograpsus testudinatus, from Shikine-jima Island in the Izu archipelago. 21, 31–36 (2019).

39. J. M. Logan et al., Lipid corrections in carbon and nitrogen stable isotope analyses: comparison of chemical extraction and modelling methods. Journal of Animal Ecology 77, 838–846 (2008).

40. E. Svensson, S. Schouten, E. C. Hopmans, J. J. Middelburg, J. S. S. Damste, Factors controlling the stable nitrogen isotopic composition (δ^15^N) of lipids in marine animals. PLoS ONE 11, (2016).

41. J. C. De Winter, Using the Student’s t-test with extremely small sample sizes. Practical Assessment, Research & Evaluation 18, (2013).

42. K. St. John Glew, L. J. Graham, R. A. R. McGill, C. N. Trueman, Spatial models of carbon, nitrogen and sulphur stable isotope distributions (isoscapes) across a shelf sea: An INLA approach. Methods in Ecology and Evolution 10, 518–531 (2019).

43. C. J. Thomas, L. B. Cahoon, Stable isotope analyses differentiate between different trophic pathways supporting rocky-reef fishes. Marine Ecology Progress Series 95, 19–19 (1993).

44. S. I. Kiyashko, T. A. Velivetskaya, A. V. Ignatiev, Sulfur, carbon, and nitrogen stable isotope ratios in soft tissues and trophic relationships of fish from the near-shore waters of the peter the great bay in the Sea of Japan. Russian Journal of Marine Biology 37, 297 (2011).

45. W. D. K. Reid et al., Spatial differences in East Scotia Ridge hydrothermal vent food webs: influences of chemistry, microbiology and predation on trophodynamics. PLOS ONE 8, e65553 (2013).

46. B. Fry, Food web atructure on Georges Bank from atable C, N, and S isotopic compositions. Limnology and Oceanography 33, 1182–1190 (1988).

47. K. L. Erickson, S. A. Macko, C. L. Van Dover, Evidence for a chemoautotrophically based food web at inactive hydrothermal vents (Manus Basin). Deep Sea Research Part II: Topical Studies in Oceanography 56, 1577–1585 (2009).

48. B. Fry, W. Ruf, H. Gest, J. M. Hayes, Sulfur isotope effects associated with oxidation of sulfide by O2 in aqueous solution. Chemical Geology: Isotope Geoscience section 73, 205–210 (1988).

